# Pan-Retinal Characterisation of Light Responses from Ganglion Cells in the Developing Mouse Retina

**DOI:** 10.1101/050393

**Authors:** Gerrit Hilgen, Sahar Pirmoradian, Daniela Pamplona, Pierre Kornprobst, Bruno Cessac, Matthias H. Hennig, Evelyne Sernagor

## Abstract

We have investigated the ontogeny of light-driven responses in mouse retinal ganglion cells (RGCs). Using a large-scale, high-density multielectrode array, we recorded from hundreds to thousands of RGCs simultaneously at pan-retinal level, including dorsal and ventral locations. Responses to different contrasts not only revealed a complex developmental profile for ON, OFF and ON-OFF RGC types, but also unveiled differences between dorsal and ventral RGCs. At eye-opening, dorsal RGCs of all types were more responsive to light, perhaps indicating an environmental priority to nest viewing for pre-weaning pups. The developmental profile of ON and OFF RGCs exhibited antagonistic behavior, with the strongest ON responses shortly after eye-opening, followed by an increase in the strength of OFF responses later on. Further, we found that with maturation receptive field (RF) center sizes decrease, responses to light get stronger, and centers become more circular while seeing differences in all of them between RGC types. These findings show that retinal functionality is not spatially homogeneous, likely reflecting ecological requirements that favour the early development of dorsal retina, and reflecting different roles in vision in the mature animal.

## Introduction

The onset of visual experience in mouse occurs around postnatal day (P) 12, at eye opening. Although the retina cannot experience patterned vision beforehand, it is remarkable that RGCs are already capable of encoding information originating from photoreceptors and transmit it to retinal central targets as soon as eyes open. However, these early light responses are far from mature, and they progressively acquire their adult features while the retina develops^1–4^. In mouse, RGC dendritic stratification in the ON and OFF layers of the inner plexiform layer matures after eye opening^5^ and light-driven activity guides the refinement of synaptic connectivity^6,7^. Consequently, RF sizes^8,9^ and complex RF properties such as direction and orientation selectivity^10,11^ keep maturing after the onset of visual experience. Yet, despite ongoing maturation after eye opening, longitudinal studies of RF properties have never been fully documented^3^.

One important yet often neglected issue is that the retina is not uniformly organised from a functional perspective. Indeed, dorsal, ventral, nasal and temporal domains have evolved to enable optimal encoding of specific features in the visual scene. For example, mouse cones co-express medium wavelength and short wavelength opsins (M-opsin and S-opsin), with a dorsal-to-ventral increasing gradient in S-opsin (and opposite for M-opsin)^12–16^. These dorso-ventral gradients affect RGC responses in adult animals with respect to their spectral tuning^17–19^, improving encoding of achromatic contrasts^20,21^ and providing evolutionary advantages for visual tasks^22^. The topographical organisation of some RGC subtypes also exhibits dorso-ventral non-uniformity^23,24^. However, nothing is known about the developmental consequences of these inhomogeneities.

Here we present a longitudinal study of RGC RF properties in the developing mouse retina from eye opening up to maturity with emphasis on dorso-ventral topographical differences. We recorded simultaneously from hundreds to thousands of RGCs at near pan-retinal level using the high-density large-scale CMOS-based Active Pixel Sensor multielectrode array (APS-MEA) featuring 4096 electrodes (42 μm pitch) arranged in a 64x64 configuration, covering an active area of 7.12 mm^2 25–27^, allowing us to discriminate topographical differences in light responses. We classified RGCs as ON, OFF and ON-OFF, measuring basic firing properties such as latency, peak amplitude and response duration at different contrast levels. We completed our study by determining the spatio-temporal properties of their RF central areas throughout development using a novel, high resolution stimulus approach.

## Results

### Simultaneous pan-retinal recording from the dorsal and the ventral retina

The spatial extent (7.12 mm^2^) of the APS-MEA chip allowed us to record simultaneously from dorsal and ventral RGCs over large retinal areas (Fig. 2A). The small electrode pitch (42 μm) enables sampling from many individual RGCs from these areas, providing us with an unbiased very large analytical sample size (see Table 1) and helps to reduce the amount of experiments/animals needed. Figure 2A shows that it is possible to visualise the outline of the retina, to quantify activity levels on individual channels and to delineate various retinal areas on the APS-MEA chip simply by looking at spiking activity. For each channel the log spike count of a full field stimulus experiment (see Methods) from a P13 (Fig. 2a) and an adult (P38, Fig. 2c) retina were pseudo color-coded and plotted according to their position on the 64⨯64 APS-MEA chip. This generates activity maps showing that the outline of the P13 retina is smaller compared to the P38 retina and that the spike rate is overall higher in all channels in that younger retina. Figures 2b and 2c illustrate responses to these same full-field stimuli in both retinas following spike sorting and RGC classification (see Methods), yielding spike rasters and histograms for dorsally and ventrally located ON (green), OFF (purple) and ON-OFF (blue) RGCs. The responses of P13 ON cells to the alternating full field stimulus were much stronger and more sustained compared to the responses of OFF and ON-OFF RGCs in the same retina (Fig. 2b) and to all responses in the P38 retina (Fig. 2d). But OFF and ON-OFF cell responses were more clearly defined at P38 than at P13. Interestingly, the ON cell responses were stronger in the dorsal than in the ventral part of the retina (Fig. 2b, compare left and right). In contrast, at P38, ON responses were somewhat stronger and briefer in the ventral part (Fig. 2d right) compared to the dorsal part, whereas it was the opposite for OFF responses (Fig. 2d left).

**Table 1.**
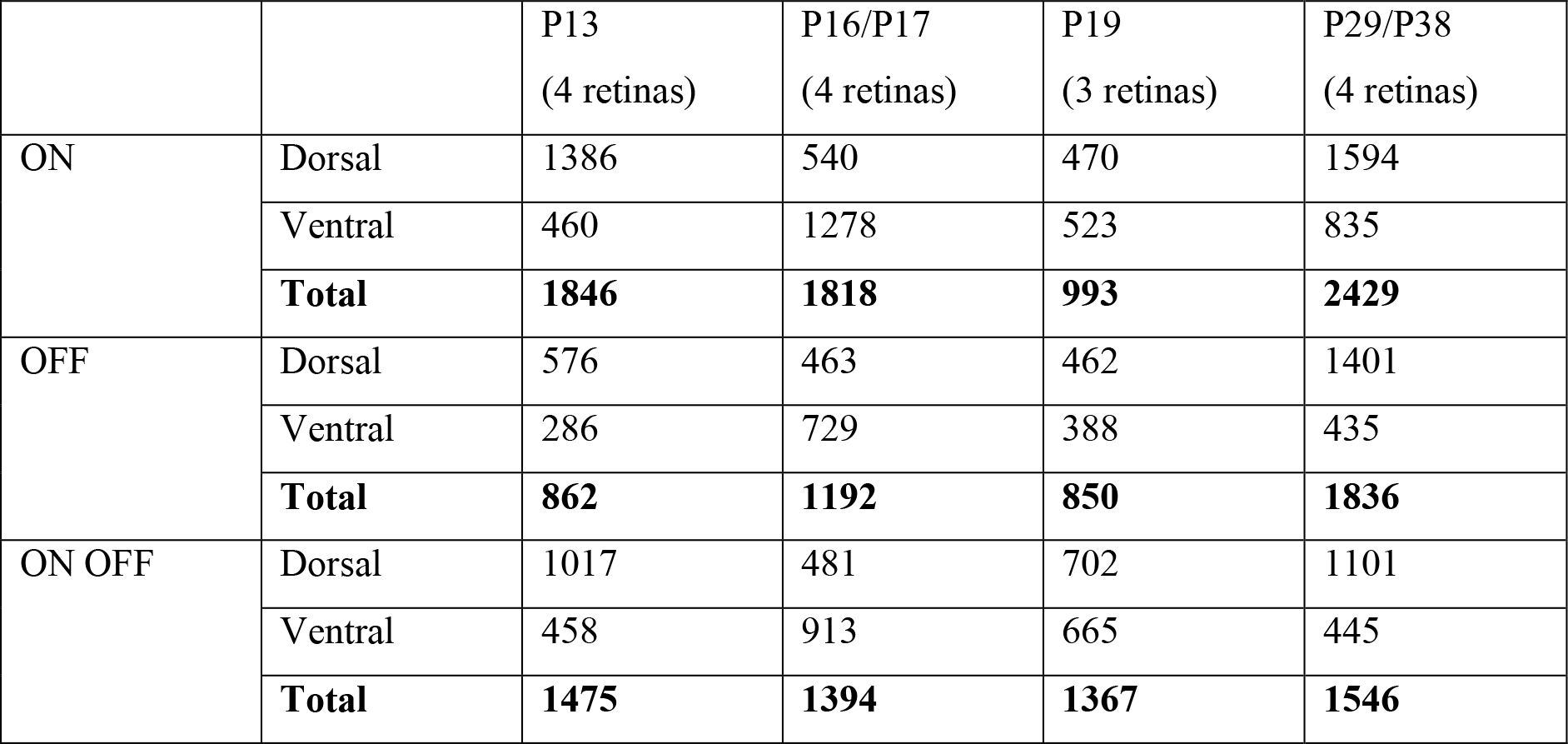
Numbers (n) of dorsal and ventral ON, OFF and ON-OFF RGCs for the different age groups used for Fig. 3–6.

|  |  | P13<br>(4 retinas) | P16/P17<br>(4 retinas) | P19<br>(3 retinas) | P29/P38<br>(4 retinas) |
| --- | --- | --- | --- | --- | --- |
| ON | Dorsal | 1386 | 540 | 470 | 1594 |
|  | Ventral | 460 | 1278 | 523 | 835 |
|  | **Total** | **1846** | **1818** | **993** | **2429** |
| OFF | Dorsal | 576 | 463 | 462 | 1401 |
|  | Ventral | 286 | 729 | 388 | 435 |
|  | **Total** | **862** | **1192** | **850** | **1836** |
| ON OFF | Dorsal | 1017 | 481 | 702 | 1101 |
|  | Ventral | 458 | 913 | 665 | 445 |
|  | **Total** | **1475** | **1394** | **1367** | **1546** |

This simple pan-retinal visualisation demonstrates that the development of basic firing properties varies not only for different RGC types but even for dorsal versus ventral ON and OFF RGCs. Next, we quantified these stimulus-driven responses for different RGC types at different ages (Fig. 3), and we examined how responses change when the full-field stimulus was presented with different Michelson contrasts (Fig. 4–6).

### Ontogeny of dorsal and ventral light response features for different RGC types

Full field stimuli were presented to retinas of different ages, and post-stimulus time histograms (PSTH) were generated for every RGC. The results were classified into 4 age groups: P13 (4 retinas, 4183 RGCs in total), P16/P17 (2 P16 and 2 P17 retinas, 4404 RGCs), P19 (3 retinas, 3210 RGCs) and adult (2 P29 and 2 P38 retinas, 5811 RGCs) and further divided into ON, OFF and ON-OFF (see Methods) dorsal and ventral RGCs (Table 1). For the results in Figures 3–6 we are referring to these groups. From individual PSTHs, we extracted the peak amplitude, time to peak and response duration for light onset and offset (see Fig. 1a). Fig. 3a summarises results for ON (left), OFF (middle) and ON-OFF (right) RGCs at all ages (x-axis), illustrating the dorsal (green) and ventral (blue) mean peak amplitude *A1* (ON cells), *A2* (OFF cells) and *A1 + A2* (ON-OFF) (see Fig. 1a). In line with examples in Fig. 2, dorsal P13 ON cells were significantly more active (Fig 3a, left) than OFF and ON-OFF cells in both dorsal and ventral areas. There is also a progressive decrease in A1 from P13 to adulthood for dorsal ON cells, whereas A2 increases with age for all OFF cells, dorsal and ventral (Fig. 3a, middle). Except for P13, A2 values were significantly higher in dorsal OFF than in ventral OFF cells. ON-OFF cells, on the other hand, did not show significant changes between the youngest and oldest age groups, and their responses were overall weaker compared to the other cell types (Fig. 3a, right).

**Figure 1.**
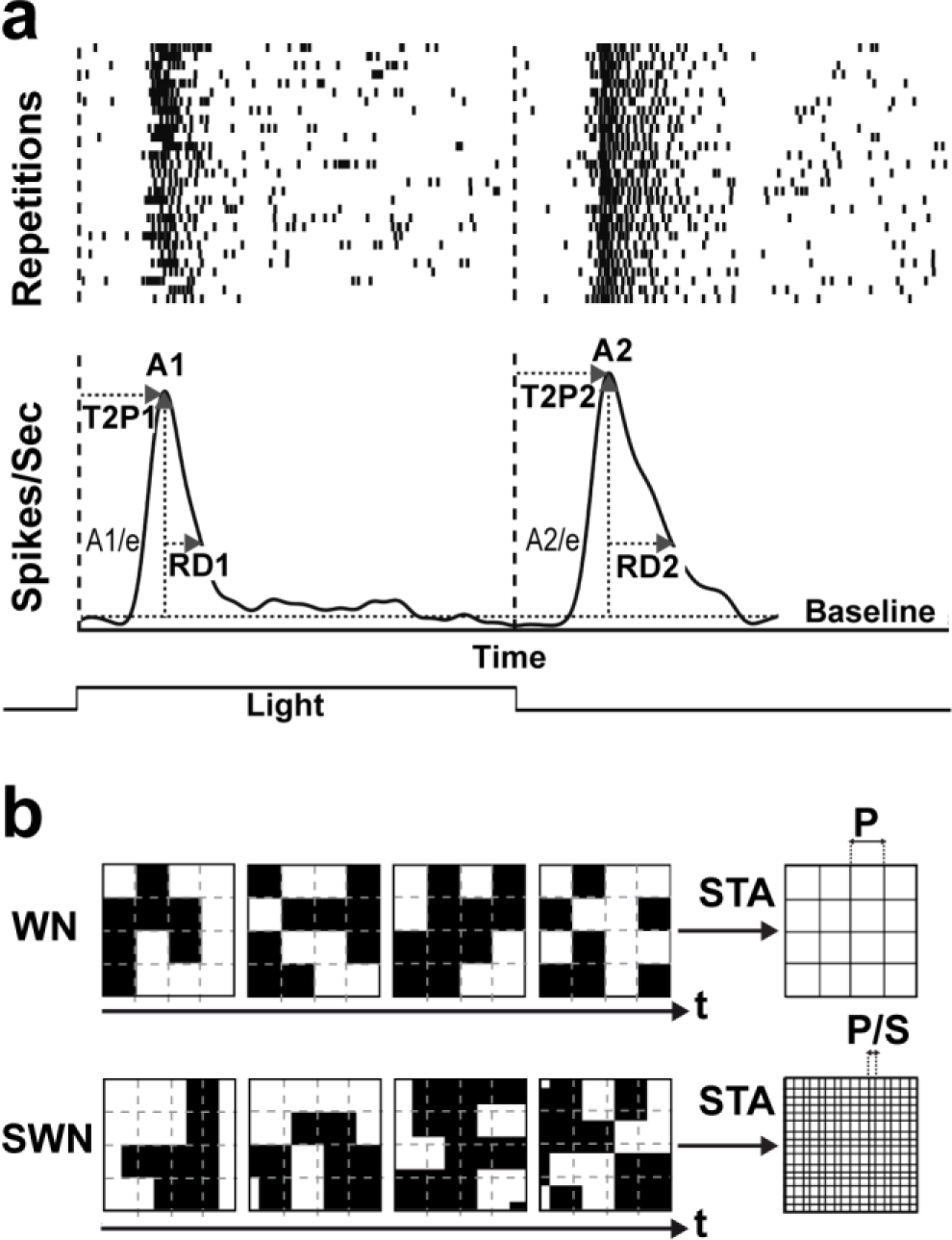
Response parameter calculations and shifted white noise paradigm. a) Top row is showing a spike raster of a full field stimulus (30 trials, 2 sec on, 2 sec off) where each dot is representing a spike, bottom row shows the averaged trials (see methods, converted to Spikes/Sec). Baseline firing rate was estimated by looking at each unit’s activity (Spikes/Sec) before stimuli were presented (>30 sec). *A1* (Spikes/Sec) is the first maximum peak (measured from each unit’s baseline) after stimulus onset and *A2* the first maximum peak after stimulus offset. *T2P1* (in ms) is the time from stimulus onset to *A1*, *T2P2* the time from stimulus offset, respectively. *RD1* (in ms) is the time, so-called response duration, after *A1* where the response is still above *A1/e. RD2* is the response duration after *A2* to *A2/e.* b) Top is showing the standard white noise paradigm with static located pixels and the resulting resolution (*P*) of the spatial STA profile. Bottom is showing the shifted white noise paradigm but with shifted (*S*) pixel location. This results in a much finer resolution *(P/S)* for the spatial STA profile.

**Figure 2.**
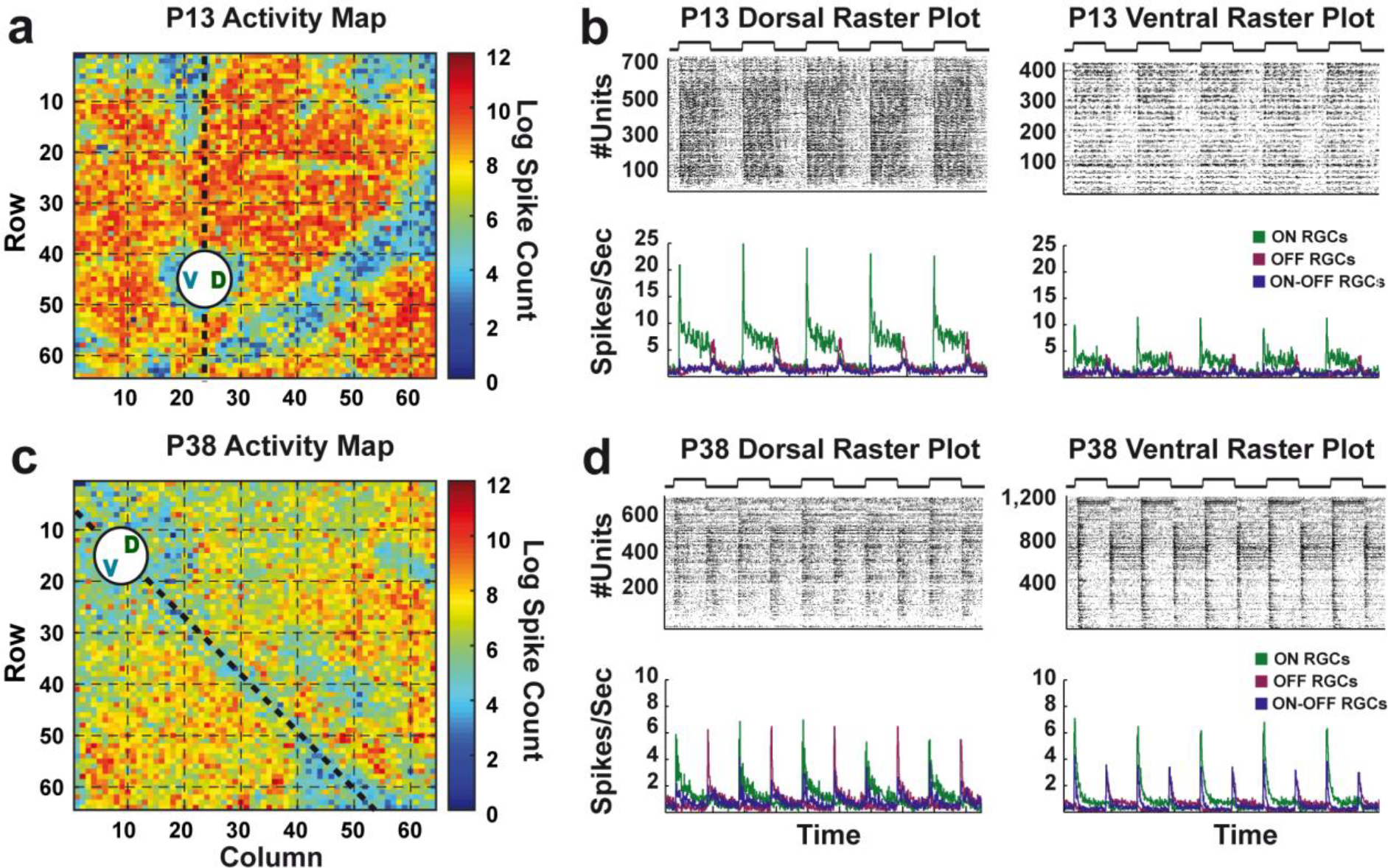
Activity maps and spike raster plots of a P13 and a P38 retina. a, c) The *Log* spike count (full field stimulus experiment) for each channel from a P13 (a) and a P38 (c) retina is pseudo color-coded and plotted according to electrode coordinates (64 ⨯ 64 array). This results in a visualisation of the retina outline and gives an overall estimation of the number of active channels. **c, d**) Spike raster plots from the same P13 (b) and P38 (d) experiment used for a) and c), respectively, but after spike sorting. Each dot is representing a spike in an alternating full field stimulus experiment and dots are color-coded: green = ON RGCs, purple = OFF RGCs, blue = ON-OFF and the raster/rate plots are divided into dorsal (b,d left) and ventral RGCs (b, d right). The binned (25 ms) average response (Spikes/Sec) of all RGCs is plotted below the raster plots.

**Figure 3.**
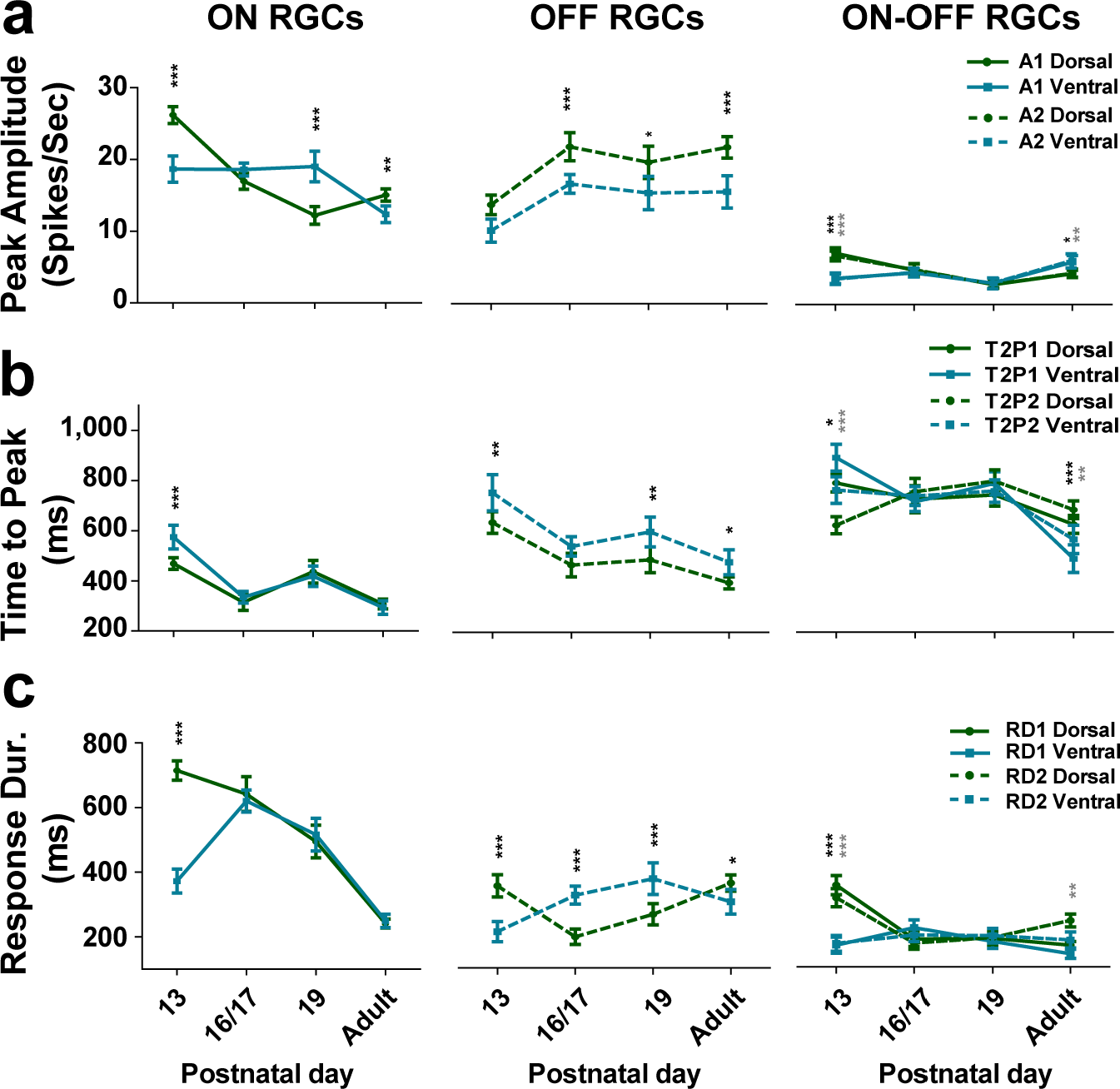
Response properties of different RGC types from P13 to adult. a, b, c: respectively illustrate peak amplitude *(A1, A2)*, time to peak *(T2P1, T2P2)* and response duration *(RD1, RD2)* for dorsal (green) and ventral (blue) ON (left), OFF (middle) and ON-OFF (right) RGCs for each age group (mean values with 95% confidence interval, n: see Table 1). For ON-OFF cells the light on-and offset values were calculated as ON-OFF 1 (solid line) and ON-OFF 2 (dotted line), respectively. Significance: * = p < 0.05; ** = p < 0.01; *** = p < 0.001. Dark asterisks are for onset values, grey asterisks for offset values, no asterisks mean not significant.

**Figure 4.**
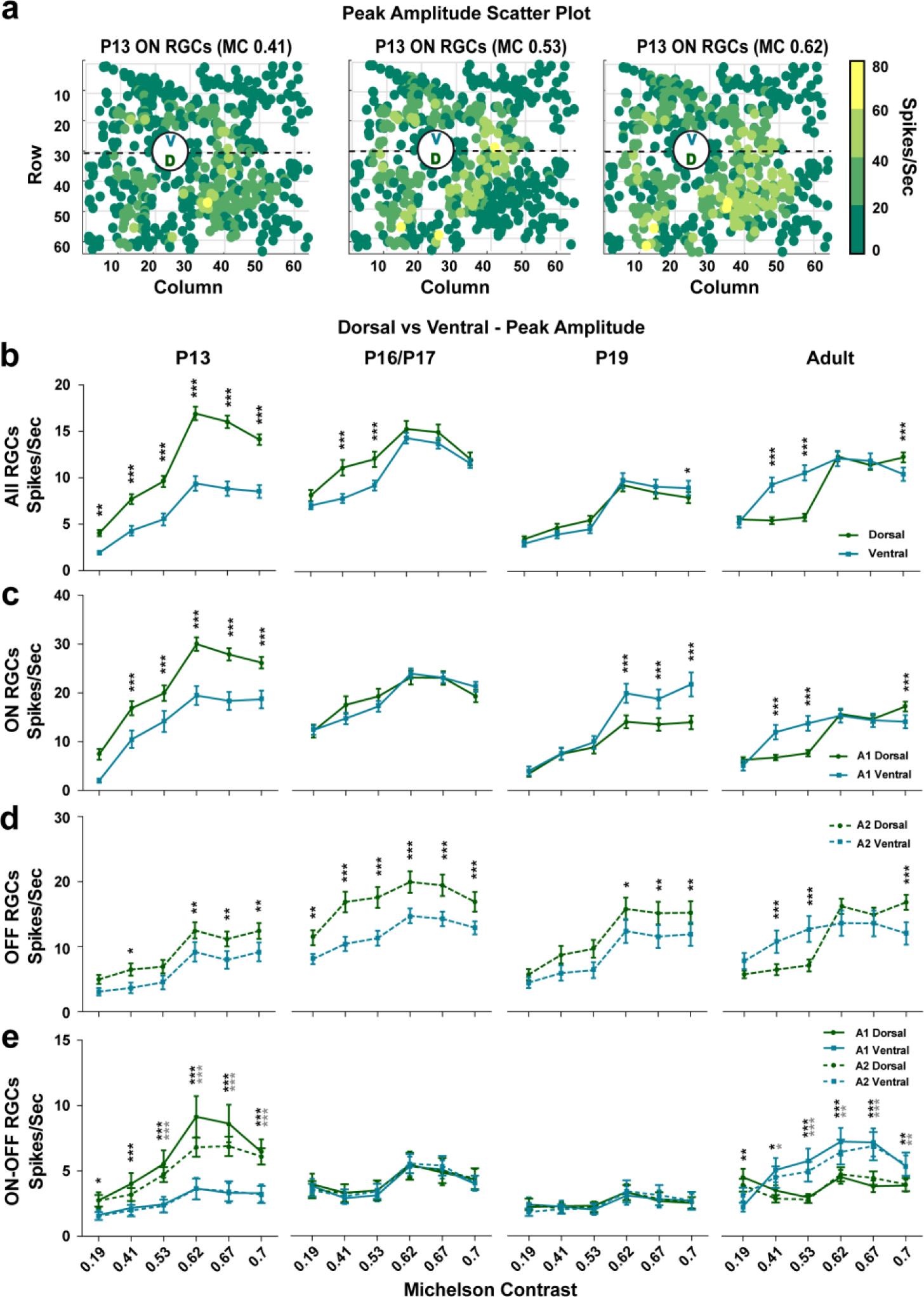
Dorsal-ventral gradient of peak amplitudes to different contrasts after eye-opening. a) ON peak amplitudes (*A1* in Spikes/Sec) for three different full field contrasts (0.41, 0.53, 0.62) from a single P13 retina are plotted in pseudo colors according to firing strength and electrode position. For visualisation, the individual x,y electrode position for the peak values was slightly randomly shifted (+/− 0.25) because after spike sorting multiple RGC units are assigned to the same electrode position. b-e) Mean peak response amplitudes to different full field Michelson contrasts (0.19, 0.41, 0.53, 0.62, 0.67) for all (b), ON (c), OFF (d) and ON-OFF (e) RGC types for all age groups (ascending from left to right) with respect to their dorsal (green) or ventral (blue) location. All plot conventions are like for Figure 3.

**Figure 5.**
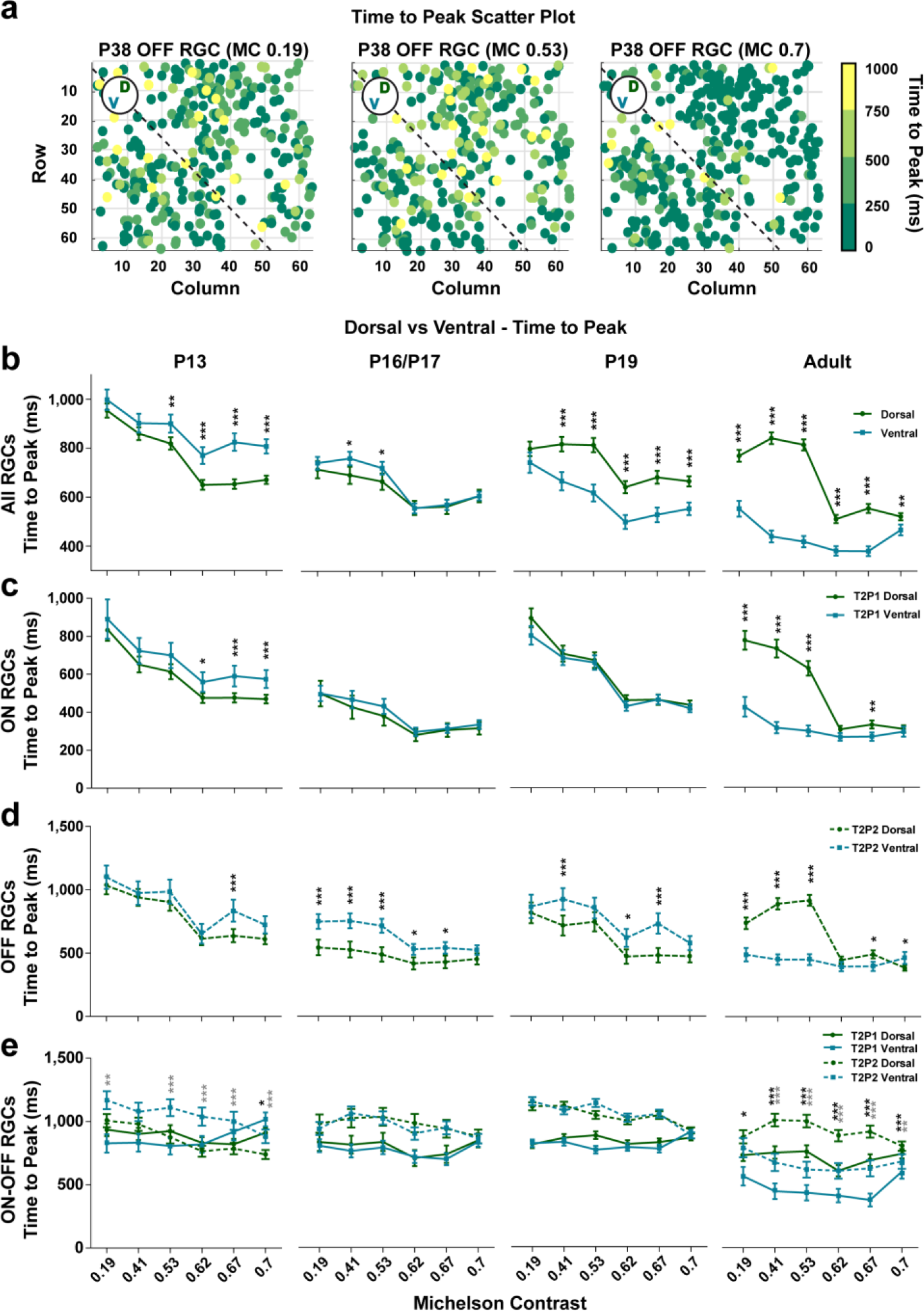
Dorsal-ventral differences for the time to peak values at lower contrast in the adult retina. a) OFF RGC color-coded response times to peak (*T2P2* in ms) from a single P38 at three different full field contrasts (0.19, 0.53, 0.7) plotted according to their electrode position. **b-e**) Mean times to peak plotted for the different full field Michelson contrasts (x axis, 0.19, 0.41, 0.53, 0.62, 0.67) for all (b), ON (c), OFF (d) and ON-OFF (e) RGC types and all age groups. All plot conventions are like for Figure 3.

**Figure 6.**
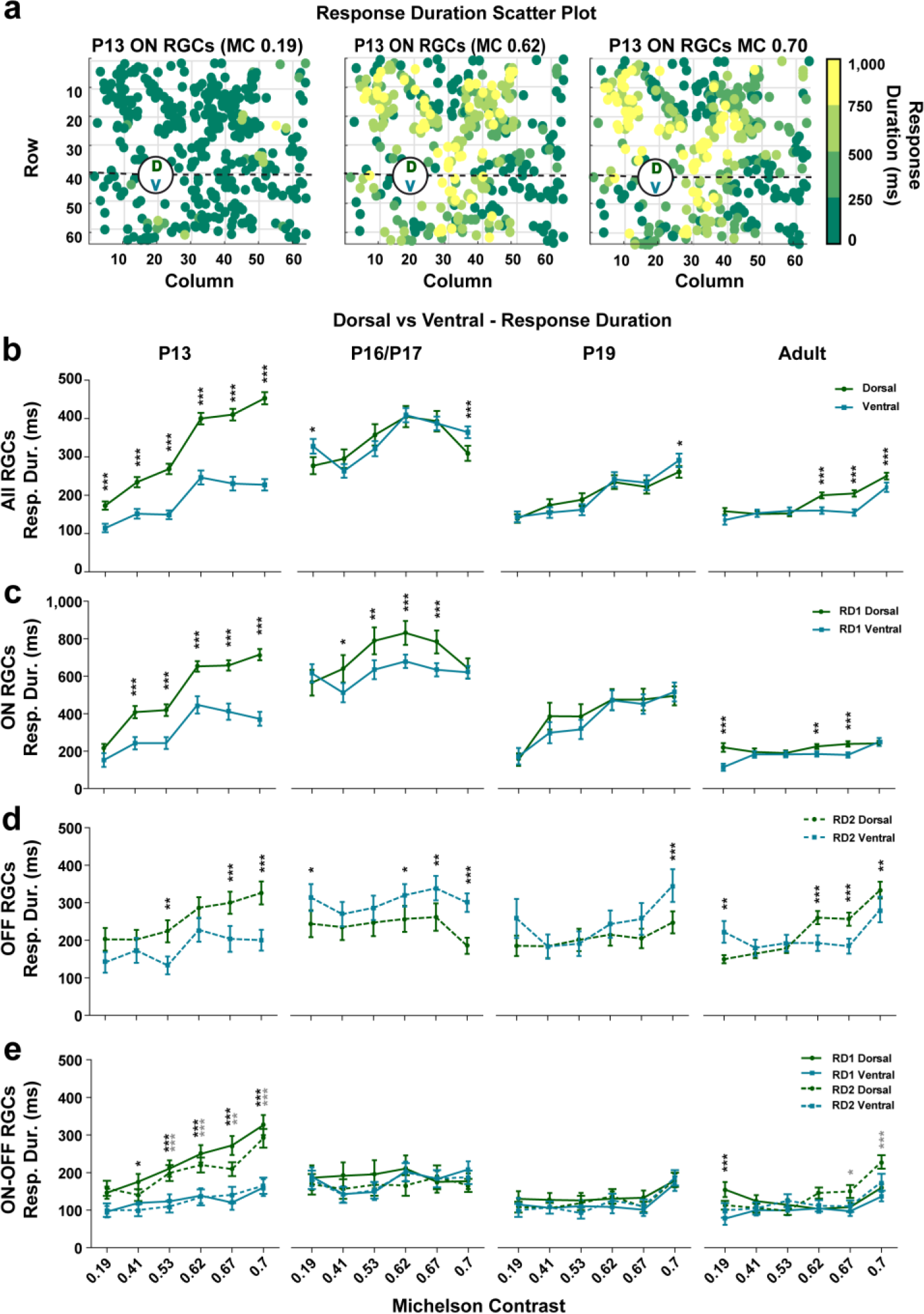
Responses to light are more sustained for all dorsal RGC types than for their ventral counterparts. a) The ON RGC response duration times (*RD1* in ms) from a single P13 retina of three different full field contrasts (0.19, 0.62, 0.7) were pseudo colour-coded plotted according to their electrode position. **b-e**) The mean response duration times to different full field Michelson contrasts (x axis, 0.19, 0.41, 0.53, 0.62, 0.67) for all (b), ON (c), OFF (d) and ON-OFF (e) RGC types and all age groups. All plot conventions are like for Figure 3.

How fast these responses to full field stimuli are at different ages was evaluated by measuring the time from stimulus on- or offset to the peak amplitude (*T2P1* and *T2P2*, respectively). We found that T2P1 in ON cells and T2P2 in OFF cells progressively decrease from P13 to adulthood, with slower responses in OFF cells (Fig. 3b, middle). ON-OFF cells had overall slower response onsets than ON and OFF cells. Overall, ventrally located RGCs exhibited slightly more sluggish responses.

How prolonged a RGC response is was defined by measuring the response duration following the peak (*RD1* and *RD2*, respectively). The spike rasters in Fig. 2b clearly show that dorsal ON cells respond in a much more sustained fashion at P13. This observation was confirmed by plotting RD1 and RD2 values (Fig. 3c, left), revealing that dorsal ON responses are significantly more sustained at P13. Subsequently, RD1 values are similar in dorsal and ventral ON cells, gradually dropping down until adulthood (Fig. 3c, left). In contrast, OFF cells become moderately more sustained between P13 and adulthood, eventually producing slightly longer adult responses than ON cells (Fig. 3c, middle). Finally, ON-OFF cells have overall shorter responses to light, and the only developmental change we observed was more prolonged responses in dorsal RGCs at P13 (Fig. 3c, right).

Taken together, dorsal light responses are more prominent after eye-opening and the peak amplitudes and response durations of ON and OFF cells showed an antagonistic behaviour from eye opening to maturity (with the most pronounced changes in ON cells) whereas the ON-OFF cell responses exhibit less developmental variability.

### Ventral and dorsal difference – peak amplitude

Next, we investigated how these developmental changes vary with the stimulus contrast. Using the same age groups as above, we recorded full field responses with different Michelson contrasts (MC, see Methods). Figure 4a illustrates the firing peak amplitudes for all ON cells (*A1*) from a P13 retina for three different full field contrasts (0.41, 0.53, 0.62) plotted with respect to their electrode position on the array. Overall, the dorsal side clearly shows stronger activity (yellow), especially for MC 0.62. To establish whether this dorsal-ventral trend is present in all retinas of the same age group and/or between the age groups, we calculated A1 and A2 for all full field contrasts for all RGC types (Fig. 4b) but also separately for ON (Fig. 4c), OFF (Fig. 4d) and ON-OFF (Fig. 4e) types. At P13, all dorsal RGC types exhibited significantly higher peak firing rates than ventral cells at all contrast levels (Fig. 4b–e left). The highest values and most pronounced differences between dorsal and ventral cells were observed at 0.62 Michelson contrast rather than at the highest contrast (MC 0.7). At P16/17 and P19, dorsal OFF RGCs were also significantly more active than their ventral counterparts (Fig. 4d middle), whereas there were no significant differences between ON-OFF cells (Fig. 4e middle). At P19, the trend reversed with ventral ON cells becoming significantly more active than their dorsal counterparts for the three strongest contrast levels (Fig. 4c middle) whereas that trend moved towards lower contrast levels in adult retinas and virtually disappeared at high contrasts (Fig. 4c, d right). Interestingly, and in contrast to P13 (Fig. 4e left), ventral adult ON-OFF cells were significantly more active (for both *A1* and *A2*) than their dorsal counterparts at all but the weakest contrast levels. Taken together, dorsal units are strikingly more active after eye-opening, an effect that disappears in older retinas.

### Ventral and dorsal difference – time to peak

Figure 5a shows *T2P2* values for all OFF cells from a mature P38 retina for three different full field contrasts (MC 0.19, 0.53, 0.7). Here, dorsal responses appear more sluggish for lower contrasts. Similar trends were observed for all adult RGC types (Fig. 5b–e right), with dorsal RGCs showing significantly longer times to peak than ventral cells, especially prominent at lower contrasts. However, at P13 all ventral RGCs had more sluggish responses at the highest MC levels (Fig. 5b–e left) whereas this was shifted towards lower contrast in OFF P16/P17 and P19 RGCs. There were no differences between dorsal and ventral ON and ON-OFF cells at P19 and P16/P17 (Fig. 5c, e middle). In summary, there is a negative developmental shift of the response time to peak for all RGC types, with the most sluggish responses found in ventral RGCs after eye-opening and slower responses in dorsal units in adults.

### Ventral and dorsal difference – response duration

Figure 6a illustrates response durations for all ON cells in a P13 retina for three different contrast levels (MC 0.19, 0.62, 0.7), with longer responses developing at higher contrasts in the dorsal part of the retina., At P13 dorsal RGCs exhibit more prolonged responses at most contrast levels (Fig. 6b–e left)., and there is a similar trend for ON cells at P16/17 (Fig. 6c, middle). In contrast, the trend was inverted for OFF cells at P16/17, with longer responses in ventral cells (Fig. 6d, middle). As for adult retinas, differences between dorsal and ventral cells became virtually non-existent, except for small differences at higher contrast levels (e.g. longer ventral OFF responses at MC 0.7 at P19; longer dorsal OFF responses MC 0.62, 0.67 and 0.7 in adult retinas).

In summary, there are significant differences between the dorsal and ventral response properties (peak firing, time to peak and response duration) for different RGC types from eye-opening to maturity. Shortly after eye opening, dorsal RGCs exhibit much stronger and more sustained responses to full field flashes than ventral cells. These dorsal-to-ventral differences fade as development progresses, and interestingly, in many cases the gradient even reverses to ventral-to-dorsal.

### Maturation of RGC RF centers

We used a novel checkerboard stimulus, shifted white noise (SWN, see Methods), to characterise the maturation of ON and OFF RGC RF centers from eye-opening to maturity. As shown in Pamplona et al.^28^, shifted white noise stimulus considerably improves the spatio-temporal resolution of the receptive field computed via STA. This allows therefore a better accuracy in receptive field characterisation. Such an accuracy is necessary to thoroughly describe the variability in receptive fields during development. The STA is the mean stimulus that precedes a spike and the cell type is inferred from the STA polarity for the phase closest to the spike time 0 (negative for OFF and positive for ON RGCs). Figure 7 shows the temporal STA signal strength (see Methods) and the spatial profile for selected ON (yellow, Fig.7a) and OFF (blue, Fig. 7b) example cells from all used age groups. The most striking developmental change we observed is that both negative and positive STA peak signal strength values are much weaker at P13–16 than later during development. Quantifying the STA strength (see Methods and legend) for all ON (Fig 7c) and OFF (Fig 7d) cells show that the STA strength significantly increases during development with the main change between P16–19. We did not observe significant differences in STA strength between dorsal and ventral ON or OFF cells at most ages, but overall, we found that responses were significantly stronger in adult RGCs.

**Figure 7.**
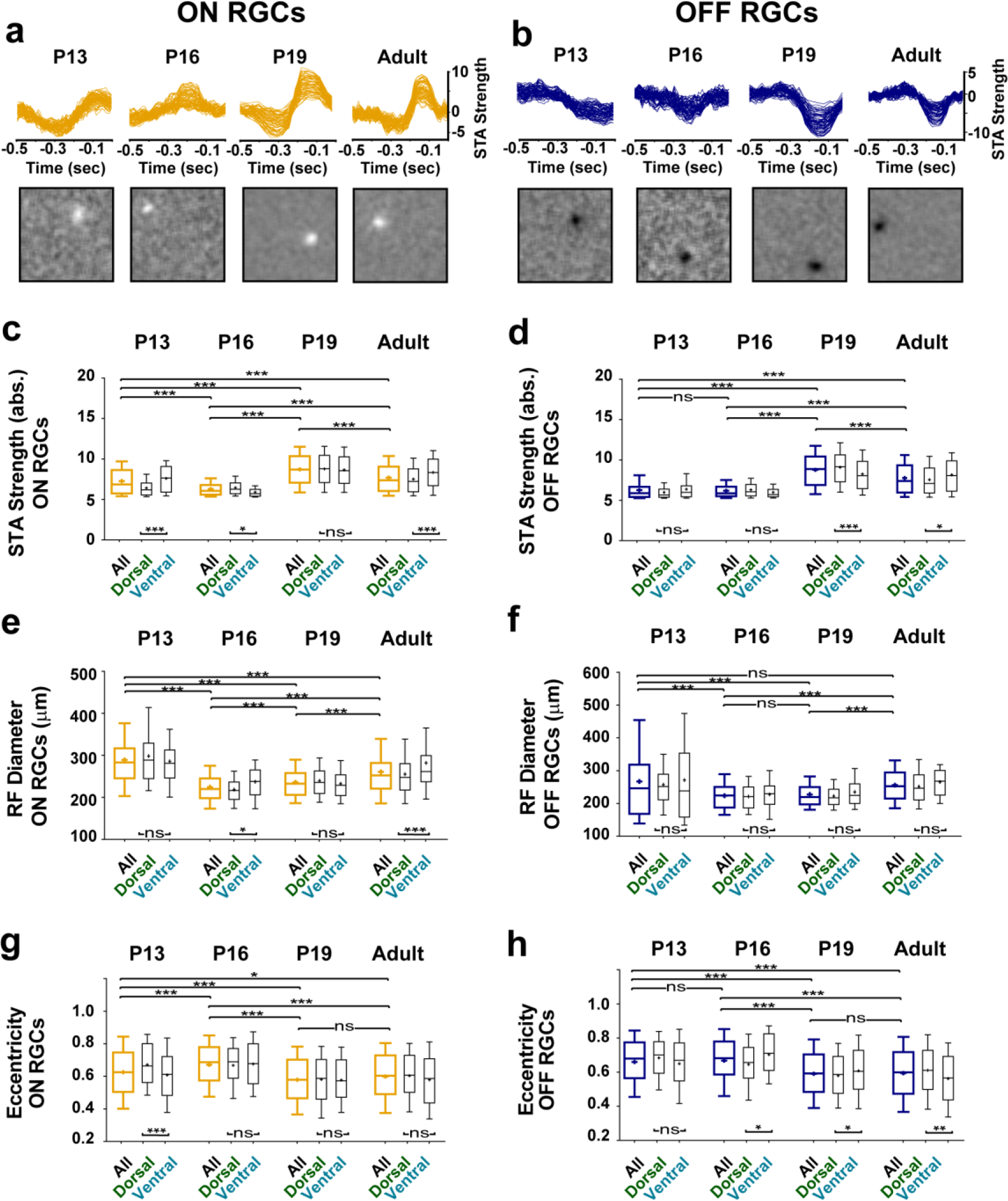
Shifted White Noise used to study the development of RGC RF central areas. **a, b**: STA signal strength (top) and spatial profiles (bottom) for selected example RF central areas from ON (a) and OFF (c) RGCs from all 4 age groups (see below). **c-h**: box plots (whiskers: 10–90 percentile, mean indicated by + symbol) of ON (yellow boxes; P13−N=2 retinas, n = 449 cells; P16−N=2, n=425; P19−N=2, n=926; adult−N=2; n=698) and OFF (blue boxes; P13−N=2, n=309; P16−N=2, n=239; P19−N=2, n=794; adult−N=2, n=514) RGCs from all retinal areas for STA signal strength (c, d), RF diameters (e, f), and RF eccentricity (g, h). All values are also shown separately for dorsal and ventral located RGCs (small black boxes). Significance: * = p < 0.05; ** = p < 0.01; *** = p < 0.001; ns = not significant.

We next used the STA to measure RF diameters (see Methods) and quantify their developmental changes from eye-opening to maturity. Figure 7e and show that RF diameters are largest at P13 both in ON (e) and OFF (f) RGCs, dropping down to a minimum at P16, followed by a marginal, nonsignificant increase at P19 and then a more significant increase from P19 to adulthood. Note that P13 ON RF diameters were significantly larger than adult RF diameters but that is not true for P13 OFF RF diameters. Since we observed significant differences in firing properties between dorsal and ventral RGCs, we also looked at RF sizes separately for these same two groups, but we found no major differences (except for adult, where ventral ON RF diameters were significantly larger than dorsal ones).

Finally, we quantified the shape of RFs by measuring their eccentricity (see Methods). As shown in Figures 7g and h, RF eccentricity for both ON and OFF RGCs significantly decreases with development; that is, the shape of RFs becomes closer to circular in later development. We did not observe significant differences between dorsal and ventral eccentricity in ON RGCs except for ventral RGCs being more circular than dorsal RGC at P13. On the opposite, dorsal and ventral RF circularity were significant differet from P16 onwards in OFF RGCs (Fig 7h, right).

## Discussion

In this comprehensive study we have shown that the basic features characterising RGC light responses have a developmental profile that depends both on cell type and retinal location. Furthermore, these properties do not develop synchronously in the dorsal and ventral retina. Finally, using a novel stimulus, we have been able to reliably characterise RGC receptive fields across development, which is difficult in young retinas where light responses to conventional white noise stimuli are weak.

We found that ON RGCs have stronger responses to light than other cell types just after eye opening. As development progresses, these responses become weaker while OFF responses gain strength (antagonistic to ON cells). The relatively high levels of spontaneous activity (residual of spontaneous waves) could not explain these results because peak amplitudes were normalised to baseline activity. Moreover, if spontaneous activity was indeed affecting our measurements, it would equally affect all RGC types, and not just ON cells. An earlier study reported that RGC light responses reach their peak strength around P28, but this study pooled ON, OFF and ON-OFF responses^7^. Pooling our data would yield similar results because of the antagonistic time course of changes in ON and OFF peak amplitudes throughout development (see Fig. 3a). The different maturation time course of ON and OFF peak firing responses may stem from differences in synaptic connectivity and membrane excitability between these cells. A previous study in mouse reported that light-evoked excitatory postsynaptic potentials are larger in ON than in OFF RGCs at P13^29^, but it also stated that both excitatory and inhibitory light-driven synaptic responses are downregulated with age, which cannot explain why we see differences between ON and OFF cells at later ages. An alternative explanation is that the expression of ionotropic glutamate receptors (AMPA/Kainate and NMDA) differs in ON and OFF RGCs. The NR2A subunit of the NMDA receptor is predominantly found at OFF synapses, while NR2B subunits are preferentially located at ON synapses in rat RGCs^30^. However, Stafford et al^31^ found no evidence for a differential localisation of the NR2B subunits at ON and OFF synapses in direction selective RGCs until P28, but no data is yet available for non-direction selective RGCs. Another explanation for our observations is that the time course of inhibitory neurotransmitter receptor expression is different for particular cell types. A good candidate is the GABAc receptor, expressed presynaptically on bipolar cell axon terminals^32^. Retinal GABAC receptor knockout results in stronger and more prolonged spiking activity^33^, similar to our observations in young ON RGCs. Further, GABAC-receptor mediated inhibition affects the regulation of glutamatergic synaptic transmission at the ON but not at the OFF bipolar-RGC synapse^34^. In contrast, Schubert et al.,^35^ stated that ON and OFF bipolar cell spontaneous inhibitory postsynaptic currents (IPSCs) are relatively similar at P12. However, the nature of light-evoked GABAergic inputs onto developing mouse bipolar cells, and how it differs for different RGC types remains to be determined (but see Discussion in^29^).

The response durations of ON and OFF RGCs (Fig. 3c) in our study showed a similar antagonistic behaviour as the peak amplitudes (Fig. 3a), but with a decrease much stronger in ON than in OFF RGCs, and a massive drop during the fourth postnatal week (Fig. 3c). In addition to the aforementioned points, GABAc receptors consist of ρ subunits and the ρ1 subunit expression peaks around P9 (start P6) and ρ2, around P15 (start P9)^36^. The difference in the timing of peak expression of these subunits may explain the differences we observe between ON and OFF RGCs. Response latencies did not exhibit conspicuous differences between different RGC types from eye opening to adulthood and the changes we observed most likely reflect ongoing activity-mediated refinement of the bipolar-RGC synapse^2,5,7,29,37,38^.

In mouse, S-opsins are co-expressed with M-opsins in cone photoreceptors and there are more S-opsins expressed in the ventral retina. S-opsin exhibit peak excitation around 360 nm^19,39^ which is in the UV spectrum. To stimulate retinas in our study, we have used a broad spectrum white light composed of red, green and blue LED lights (∼420 − 660 nm) but not UV light. Therefore it is highly unlikely that we were able to stimulate S-opsin properly. Under these conditions, dorsal RGCs have an advantage because our white light source is biased towards activation of M-opsin, which is more prevalent in the dorsal retina. Yet, our results do not demonstrate a general increase of response strength in the dorsal side *per se*, but reveal a differential dorsal/ventral development pattern different RGC types through maturation. Even though a bias in M-opsin activation could potentially explain the stronger responses we recorded at P13, it cannot be responsible for the results we found at later developmental stages, including a switch to stronger ventral responses at low contrast levels in the adult retina. S- and M-opsins are expressed before eye-opening, respectively from ∼P1 and ∼P8^13,14^. Therefore we can reasonably assume that both S- and M-opsin gradients are already fully established shortly after eye opening, when we sampled the earliest light responses, and differences in opsin expression are unlikely to explain our findings. Further, the light intensity (mean luminance 11 cd/m^2^) in our experiments was set to co-activate rods and cones. Therefore rod-mediated (by rhodopsin in the visible spectrum) responses are not saturated in our recordings^40^, and they probably reflect a large proportion of our recordings because rods outnumber cones by 35:1 in the mouse retina^41^. Stronger responses in the dorsal retina at the onset of visual experience is not an unreasonable possibility from an ecological and evolutionary point of view because at that age, pups are still gathered in the nest, near the mother, with no ecological need to look skywards, but they rather concentrate on the nest scenery, using dorsal retinal vision. In the ensuing week, retinal circuits keep developing and refining, leading to maturation of the ventral circuitry as well.

Presenting white noise checkerboard images and performing a post hoc reverse correlation is now a standard approach to estimate RGC RF centers^42,43^. However, the technique is challenging in very young retinas because of high levels of spontaneous activity and the weak sensitivity to light in young RGCs immediately after eye opening. The problem can be alleviated by increasing the number of trials and/or the pixel size without compromising the minimum resolution for reliable RF estimation^8,9,44^. Here we applied a novel approach, the SWN, which uses large checkerboard pixels (hence evoking strong responses) randomly shifted in space and time by a fraction of the pixel size, yielding better resolution. SWN images are uncorrelated across time, and the bias introduced by the correlation in space due to finite pixel size is negligible. Using this approach, we have shown that STA signal strength defined by the z-score (see Methods) peaks at P19 for ON and OFF RGCs which is in line with Cantrell et al^8^, who showed a signal strength peak at P18. The significantly higher signal strength in adult ventral RGCs is consistent for ON and OFF RGCs but antagonistic to our full field results. Further studies are thus needed to investigate the spatial response inhomogeneities in adult retinas. The main developmental increase in STA signal strength peak correlates well with the end of the programmed cell death period for bipolar cells and outer rods^45^, suggesting that the synaptic connectivity required for mature RF center responses is complete at that time, and only minor synaptic refinements occur later on, resulting in further RF expansion. We found that immature (P13) RF diameters in ON RGCs are significantly larger than later in development, while for OFF cells, RFs are similar in size at P13 and in adults, with a temporary drop during the third postnatal week. Previous studies which used conventional checkerboard images present a different developmental picture. Koehler et al.^9^ showed that ON and OFF RFs are smaller in adults than immediately after eye-opening whereas Cantrell et al.^8^ stated that OFF, but not ON RFs expand between P15–18 and that there is a significant difference for ON, but not for OFF cells between P18–25. This demonstrates that different checker board approaches can yield different results and will be elaborated in another study^28^. Additional factors such as mean luminance, adaptational state and recording length may account for these differences, and results should therefore be interpreted with caution. From our measurements of RF eccentricity, we found that RFs become more circular between P16–19. These changes are probably caused by the same factors responsible for developmental changes in STA signal strength. Interestingly this goes in line with a previous study in turtle where the authors could show that anisotropic properties of spatiotemporal RFs decrease with maturation^3^. The significant difference between dorsal and ventral eccentricity in ON P13 needs further investigation but it seems that higher STA signal strength predicts more isotropic RF centers (data not shown but see dorsal and ventral boxes in Fig 7c, d and Fig 7g, h).

In summary, using a large-scale, high-density multielectrode array, we have investigated the ontogeny of light responses in RGCs in the developing mouse retina. We were able to record from hundreds to thousands of RGCs simultaneously at pan-retinal level, and found that the refinement of RF properties strikingly differs between RGC types and between the same RGC types in the dorsal and ventral retina. These findings suggest that retinal functionality is not spatially uniform and that there might be an ecological advantage to favouring the development of dorsal light responses before the rest of the retina reaches functional maturity.

## Materials & Methods

### Retina Preparation

Experimental procedures were approved by the UK Home Office, Animals (Scientific procedures) Act 1986. Male and female wild-type mice (C57bl/6), housed under a 12 hour light-dark cycle and aged between postnatal days (P) 13–63 were used for the experiments. Mice were dark-adapted overnight and killed by cervical dislocation. Eyes were enucleated, and following removal of the cornea, lens, and vitreous body, they were placed in artificial cerebrospinal fluid (aCSF) containing the following (in mM): 118 NaCl, 25 NaHCO3, 1 NaH PO4, 3 KCl, 1 MgCh, 2 CaCh, 10 glucose, and 0.5 l-Glutamine, equilibrated with 95% O2 and 5% CO2. The ventral and dorsal orientation was marked after enucleation and confirmed by using vascular landmarks in the retina ^46^ The retina was isolated from the eye cup and flattened for MEA recordings and the ventral-dorsal, nasal-temporal orientation was noted down. All procedures were performed in dim red light and the room was maintained in darkness throughout the experiment.

### APS-MEA recordings

The isolated retina was placed, RGC layer facing down, onto the APS-MEA and flattened by placing a small piece of translucent polyester membrane filter (Sterlitech Corp., Kent, WA, USA) on the retina followed by a home-made anchor. Similarly as described elsewhere^26,27^, throughout recording, retinas were maintained at 33°C using an in-line heater (Warner Instruments LLC, Hamden, CT, USA) and continuously perfused using a peristaltic pump (∼1 ml min^−1^). Pan-retinal recordings were performed on the BioCam4096 platform with BioChips 4096S+ (3Brain GmbH, Lanquart, Switzerland), integrating 4096 square microelectrodes (21 ⨯ 21 μm, pitch 42 *μ*m) on an active area of 2.67 ⨯ 2.67 mm. The platform records at a sampling rate of 7.1 kHz/electrode when using the full 64⨯64 array and recordings were stored at 12 bits resolution per channel with a 8 kHz low-pass filter/0.8 Khz high-pass filter using 3Brain’s BrainWave software application. Data management and analysis for large-scale, high-density APS-MEA recording is particular difficult (as reviewed in^47^). To reliably extract spikes from the raw traces we used a quantile-based event detection ^26,48^ and single-unit spikes were sorted using the T-Distribution Expectation-Maximisation algorithm in Offline Sorter (Plexon Inc, Dallas, USA). Sorted units that exhibited at least >0.1 spikes/sec on average over the entire recording session were then verified by visual inspection of the detected clusters in the 2/3D principal component feature space, the calculated cluster interspike intervals with respect to the refractory period and the shape of the spike waveforms in the Offline Sorter GUI. Due to the high density of the electrodes, the same units were sometimes detected on multiple neighbouring channels. These redundant units were removed by comparing coincident spikes between neighbouring units. Briefly, for each unit, spikes occurring within +−2 frames (1 Frame = 1/7.06 ms) were detected in all units on the four closest electrodes and marked. This was done for all units, and units with more than 5% coincident spikes were iteratively removed such that for each coincident group only the one with the largest spike count was retained. On average we finally record from multiple hundreds to thousands individual RGCs in one experiment.

### Light stimuli

Light stimuli were projected onto the retina as described previously^27,47^ by exploiting the temporal and spatial resolution of the experimental platform and were attenuated using neutral density filters to high mesopic light levels (mean luminance 11 cd/m^2^).

A full field stimulus that switched from light to dark (0.5 Hz, 30 repetitions) was used to define peak response latency, duration and relative amplitude of ON and OFF responses (see Fig. 1A). The luminance contrast for this stimulus measured as the Michelson contrast was defined as (*I*_max_ − *I*_min_)/(*I*_max_ + *I*_min_) where *I*_max_ and *I*_min_ are respectively the maximum and minimum luminance and had a maximal value of 0.70. We also used full field stimuli with a series of increasing Michelson contrasts (0.19, 0.41, 0. 53, 0.62, 0.67). We estimated each unit’s instantaneous firing rate for the different full field intensities by convolving its spike train with a Gaussian kernel smoothing function (standard deviation = 25ms). We then averaged the trials (Fig. 1A) and extracted several features like the amplitude of ON and OFF responses (*A1, A2*), the time from stimulus onset or offset to peak of these response (*T2P1, T2P2* respectively) and the response duration (*RD1, RD2*). Statistical significance was evaluated using Oneway ANOVA with a Bonferroni post-hoc test (Prism, GraphPad, CA).

To classify RGCs according to their main response polarity, we measured the relative amplitude of ON and OFF responses and calculated the *Bias Index (BI)* defined as (*A1 − A2*)/(*A1 + A2*) (Carcieri et al., 2003). We used the *BI* to classify the cells into OFF (*BI* −1 to −0.33), ON-OFF (*BI* −0.33 to 0.33) and ON cells (*BI* 0.33 to 1).

### Shifted white noise

Checkerboard stimuli are routinely used to measure RF areas^42,43^. Finer resolution can be achieved using smaller unitary checkerboard pixels, but at the same time very small checkerboard pixels may not be able to elicit reliable and repeatable responses, if at all, necessitating to reach a trade-off. Here we use an improved checkerboard stimulus, so-called shifted white noise (SWN), where checkerboard pixels are shifted randomly in space at fixed time steps^28^. With this novel approach, the checkerboard pixel size is large enough to reliably evoke significant responses, but at the same time, the RF resolution can be very fine, given by the shift size (Fig. 1B). The SWN at position *x, y* and time *t* is defined as:

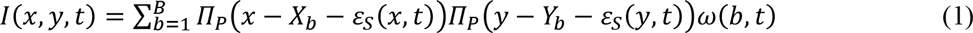

Images are composed of *B* checkerboard pixels with a fixed size *P*, the area of each pixel is defined by a rectangular function *Πp, b* is the block index and its top left corner coordinates are *(Xb, Yb)*. The index of each pixel is given by the random variable *ω(b,t)* which is taken from a Bernoulli distribution of values −1 (color black) and 1 (color white) with equal probability 0.5 for each block *b* at each time stamp *t*. Each image is randomly shifted horizontally (shift *ɛ_x_(t)*) and vertically (shift *ɛ_y_(t)*). The shifts *ɛ_x_(t)* and *ɛ_y_(t)* are random variables taking *S* possible values with a probability *1/S*. Here we take S=4. They are redrawn at each time step. We used 17 ⨯ 17 pixels (*B* = 289) with an edge size of 160 μm (*P* = 160) and a shift step size of P/4 (40 μm) and we considered 4 steps (*S* = 0, 40, 80, 120) both for horizontal and vertical shifts. As shown in Pamplona et al, this form of stimulus considerably improves the spatial and temporal resolution of the STA^28^. The Michelson contrast was 0.7 with the same mean luminance stated before and SWN images were presented for 33 ms each (30 Hz, ∼45 min, ∼15 min for adult retinas). The spike triggered average (STA)^42^ was calculated by computing the average stimulus 500 ms (corresponding to 15 checkerboard frames) before a spike occurred. At the time point of the positive or negative peak maximum of the average temporal STA, a two-dimensional Gaussian was fitted to the corresponding spatial profile frame and an ellipse was drawn around the center with 1 SD of the Gaussian fit. The RF diameter was defined as the diameter of a circle with the same area as the ellipse (2*radius = 2 SD). We also measured the eccentricity of the fitted ellipse to see how ‘out of round’ an RF is. The eccentricity *e* is given by:

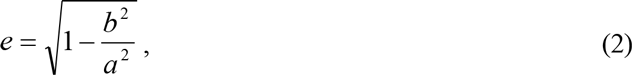

where *a* is the length of the semi-major axis (half of the longest diameter of an ellipse) and *b* is the length of the semi-minor axis (half of the shortest diameter of an ellipse). The eccentricity changes from 0 to 1, where the eccentricity of a circle is zero.

To measure STA strength at the time of the positive or negative peak maximum of the average temporal STA, we computed the z-score of STA peak amplitudes; that is for a given cell the mean of STA amplitudes was subtracted from the peak amplitude and divided by their standard deviation.

## Acknowledgements

The research received financial support from the 7th Framework Program for Research of the European Commission (Grant agreement no 600847: RENVISION, project of the Future and Emerging Technologies (FET) program Neuro-bio-inspired systems (NBIS) FET-Proactive Initiative)). We thank the IIT-NetS3 laboratory (member of the RENVISION consortium) for providing the light-stimulation system integrated with the CMOS-MEA platform used in this work.

## Author Contributions

Conceived and designed the experiments: G.H., E.S., D.P., and P.K. Performed the experiments: G.H. Analysed the data: G.H and S.P. Contributed reagents/materials/analysis tools: S.P. and M.H.H. Contributed to the writing of the manuscript: G.H., S.P., D.P., P.K., B.C., M.H.H., E.S.

